# Fungi are colder than their surroundings

**DOI:** 10.1101/2020.05.09.085969

**Authors:** Radames JB Cordero, Ellie Rose Mattoon, Arturo Casadevall

## Abstract

Fungi play essential roles in global ecology and economy, but their thermal biology is widely unknown. Mushrooms were previously noticed to be colder than surrounding air via evaporative cooling or evapotranspiration. Here we applied infrared imaging to reveal that not just mushrooms, but also molds and yeasts maintain colder temperatures than their surroundings via evapotranspiration. On average, fungal specimens are ~2.5 °C colder than the surrounding temperature. The relatively cold temperature of mushrooms can be observed throughout the whole fruiting process and at the level of mycelium. The mushroom’s hymenium appeared the coldest and different areas of the *Pleurotus ostreatus* mushroom appear to dissipate heat differently. Evapotranspiration in yeast and mold biofilms can be measured from the accumulation of condensed water droplets above biofilms; which is significantly higher than the surrounding agar. We also present a mushroom-based air-cooling system (MycoCooler™) capable of passively reducing the temperature of a closed compartment by approximately 10 °C in 25 minutes. These findings suggest that the fungal kingdom is characteristically cold. Since fungi make up ~2% of Earth biomass, their evapotranspiration may contribute to planetary temperatures in local environments. This study present new research avenues in fungal biology, biomedicine, microclimate, and sustainable energy.

## Background

Temperature controls the growth, reproduction, and ecological distribution of all life forms. The temperature of an organism depends on the balance between gaining and dissipating heat as it is influenced by its total environment (i.e., physical-chemical, biotic-abiotic, micro-macro dimensions) [1]. In theory, if the organism gains more thermal energy than it dissipates, it becomes warmer. If more thermal energy is lost, the organism may reach colder temperatures than its surroundings. When the organism and the environment each have the same temperature, there is no heat flow; hence the organism is in thermal equilibrium. Living organisms are considered dissipative systems that exist far from thermodynamic equilibrium [2]; that could mean they are warmer or colder, but not equal in temperature to their surroundings.

The temperature of microorganisms at the cellular and community level (colony or biofilm) and the mechanisms of heat exchange with their surroundings are largely unknown. Thermal biology is more advanced in the animal kingdom, where organisms can be classified based on their primary source of heat (endotherm/ectotherm) and on their capacity to maintain their body temperatures relative to their environment (homeotherm/poikilotherm) [3]. The primary source of body heat in endotherms is internal metabolism. Endothermic organisms (alias ‘warm-blooded’), like birds and mammals, can maintain relatively constant internal temperatures that range from 36 to 40 °C, regardless of any fluctuations in outside temperature. The ability to maintain stable body temperatures regardless of the primary source of heat is known as homeothermy. Most life forms are, however, ectothermic (alias ‘cold-blooded’) as their primary source of body heat is the environment. Ectotherms tend to be poikilotherms since their internal temperatures fluctuate with changes in external temperatures. Although these terms are sometimes deemed exclusive to animals, other kingdoms of life can undergo more primitive forms of thermoregulation based on similar principles.

Plants makeup ~80% of the Earth’s biomass [4] and are frequently found in environments subject to wide temperature variations and, unlike reptiles or fish, cannot displace to more favorable environments as needed [5]. They can thus be considered the “epitome of poikilothermy” [6]. On the other hand, there is evidence that some plant leaves can maintain relatively constant temperatures [7] and regulate heat loss via behavioral and physiological mechanisms. One such physiological mechanism involves the evaporation of water at their leaves’ surfaces known as evaporative cooling, transpiration, or evapotranspiration. The evaporation of water is an endothermic process that consumes thermal energy to break hydrogen bonds when water goes from liquid to a gas [8]. Stomas, microscopic pores in some leaves, regulate the water transpiration process by opening and closing in response to stimuli. Depending on the thermal environmental conditions, leaves may be warmer than air temperatures [7] or can dissipate heat via evapotranspiration, thus becoming colder than surrounding air temperature [9–11]. Consequently, plants cool themselves and their surrounding environment by evaporating water, which collectively contributes to cloud formation. Evaporative cooling is also observed in animals, where cellular structures such as sweat glands regulate the water evapotranspiration process.

Fungal, protist, archaeal, and bacterial communities are assumed to have ectothermic (that is, dependent on external sources of heat) properties considering their relatively simpler physiology and small size (or high surface area to volume ratio) [12]. However, the identification of heat-producing bacteria [13] suggests that a microbial community can produce enough thermal energy and maintain warmer temperatures than the surroundings, suggesting that microbes can exhibit endothermic (internal heat production) properties. This paper aims to evaluate the temperature of fungi in relation to their surroundings.

In geologic history, fungi pioneered the colonization of land. Today, they play a central role in balancing Earth’s ecology by breaking down decaying biological matter and providing nutrients for new growth. Surviving almost anywhere, fungal organisms can be a source of food, medicines, and a variety of biomaterials [14]. The fungal kingdom also includes species that are mutualists such as mycorrhizal fungi, as well as, species in the form of molds and yeasts that are pathogenic to animal and plant flora, causing severe public health and agricultural problems [15]. Pilei are often convex but can also form other shapes during development and between species. The area underneath the pilei, the hymenium, consists of lamellae gills or porous surfaces bearing spores. The lamellar constitution of the hymenium can increase the surface area of mushrooms by 20 folds [16]. The structural organization of the hymenium is important for spore production and release.

Previous studies and observations have noted that mushrooms, the reproductive structure of fungal mycelium, tend to be colder than their surroundings [17–20]. The first study inserted thermocouple detectors into mushrooms and suggested that the relatively cold temperatures were mediated by evaporative cooling [19]. Quantitative data of mushroom transpiration was provided in subsequent studies [17,18]. Mahajan et al. quantified the transpiration rate of *A. bisporous* whole mushrooms and developed a mathematical model to link mushroom water loss with ambient temperature and relative humidity [17]. Subsequently, Dressaire et al. quantified the rate of water loss from mushroom pilei, which can be higher than plants and enough to cool the surrounding air by several degrees Celsius [18]. In addition to thermocouple detectors, the relative cold structure of mushrooms was also noted using video thermography [20]. In this study, we applied infrared imaging to measure the temperature of wild mushrooms in their natural habitat, as well as, laboratory-grown colonies and biofilms of molds and yeasts, thus extending the concept of fungal hypothermia to the microorganism domain. We show that all these fungal life-forms appear colder than their surroundings and use the process of evapotranspiration to give off heat. Relative coldness appears to be a general characteristic observed across different fungal lifestyles, from macroscopic to microscopic fungi.

## Results

### Mushrooms, yeasts, and molds maintain colder temperatures than their surroundings

Thermal imaging of 21 mushroom species in their natural habitat and while attached to their natural substrates revealed that each was colder than their surroundings (**Fig. 1 a-f** and **Table S1**). The temperature of mushroom stalks (or stipe) recorded for some wild specimens was similar to the pilei (or cap). Yeast colonies and mold biofilms of *Candida* spp., *Cladosporium* spp., *Penicillium* spp., and *Rhodotorula mucilagenosa* also exhibited lower temperatures than the surrounding agar media following 1 h incubation at 45 °C (**Fig. 1 g-j**). The temperature differences between the fungus and its surroundings averaged ~2.5 °C and varied from ~0.5 to 5.0 °C, depending on the fungal specimen (**Fig. 1 k**). The temperatures of all fungal specimens, including those grown in the laboratory or found in their natural environment, correlated linearly with surrounding temperatures at a slope of 1 and x, y-intercepts of ~2 °C (**Fig S1**). We also found one instance where a darkly pigmented *Boletus* spp. wild mushroom pilei in natural and open-habitat conditions appeared warmer than its immediate surroundings, but relatively cold when imaged indoors, hours after detachment from its natural substrate (**Fig S2**). This observation indicates that mushrooms are relatively cold but can also warm up in open-habitat conditions via pigment-mediated heat capture from radiation energy.

**Figure 1.**
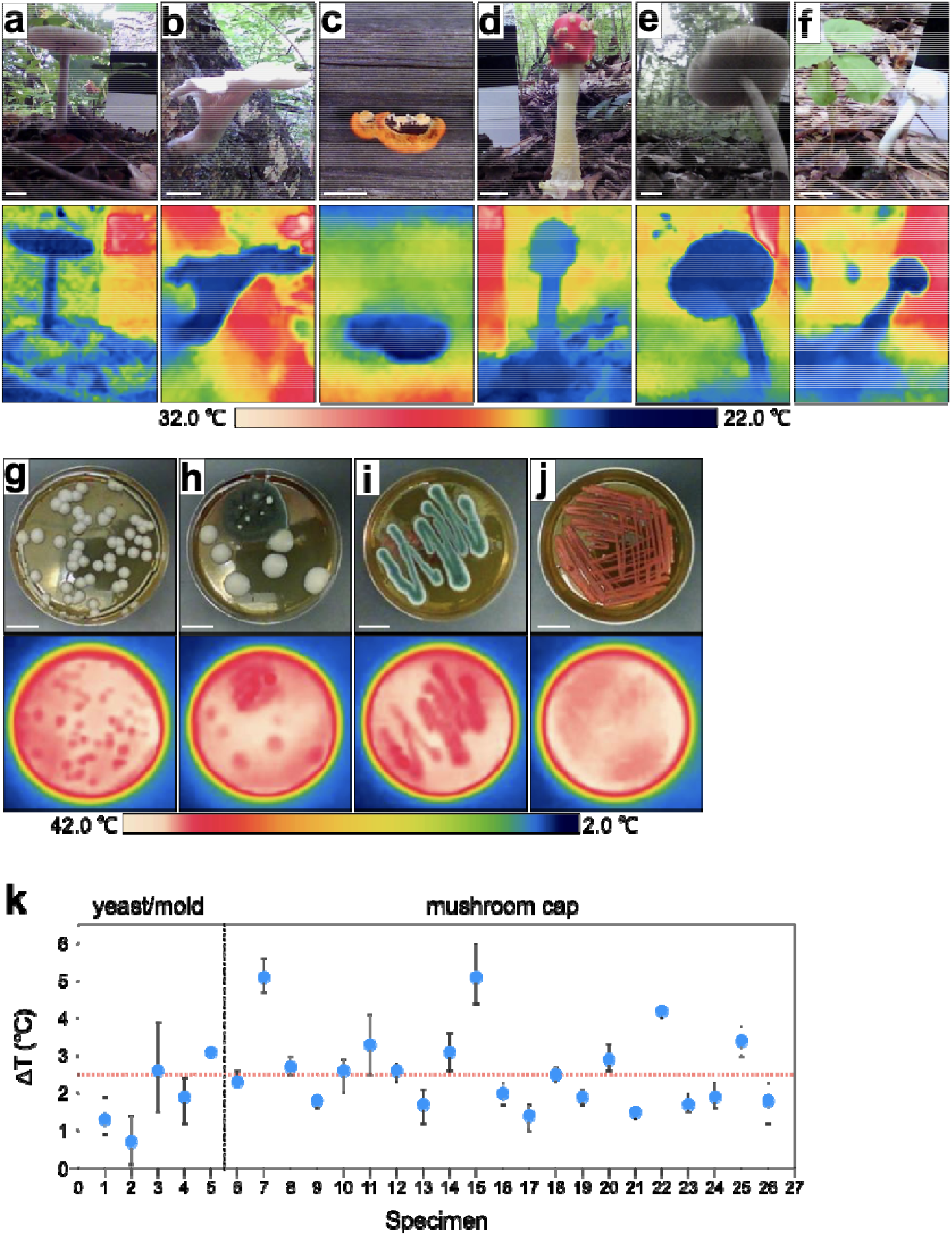
Yeasts, molds, and mushrooms are colder than their environment. Thermographic examples of wild mushrooms imaged in their natural habitat and attached to their natural substrate: (**a**) *Amanita* spp., (**b**) *Pleurotus ostreatus*, (**c**) *Pycnoporus* spp., (**d**) *Amanita muscaria* (**e**) *Amanita brunnescens,* (**f**) *Russula* spp. (**g**) yeast *Candida* spp. (also seen in **b** as white colonies), (**h**) mold *Cladosporium sphaerospermum* (dark colony), (**i**) mold *Penicillium* spp., and (**j**) yeast *Rhodotorula mucilaginosa*. (**k**) The temperature difference between the surrounding/ambient and fungal specimen. Error bars represent standard deviation. Y-intercept line indicates the mean temperature difference. The temperature values of all fungal specimens and surroundings are listed in **Table S1**.

### Change in mushroom temperature during fruiting, heating, and cooling

*Pleurotus ostreatus* grown in the laboratory at 25 °C revealed relative coldness throughout the whole fruiting process (**Fig. 2 a**). We recorded colder temperatures over time, as the mushroom flush grew in size while attached to the substrate. The mushroom flush remained relatively cold after detachment, although it was still several degrees warmer than it was when attached to the substrate. The hymenium or area underneath the pileus appeared colder than the frontal side of *P. ostreatus* pilei or stalk (**Fig. 2 b**). Notably, the fruiting site of the mycelium also remained relatively cold after mushroom detachment, approximately 2.5 °C cooler than the rest (**Fig. 2 b**). The relatively cold temperature of the *P. ostreatus* mushroom was maintained during heating, increasing from approximately 19 to 27 °C following 137 minutes incubation at 37 °C (<10 % relative humidity, RH) (**Fig. 2 c**). After heating, the mushroom flush was incubated at 4 °C (<10 % RH), and its temperature dropped from 24 to 18 °C after 104 minutes (**Fig. 2 d**). A comparison of the thermal images of the mushroom flush during heating and cooling incubations showed that different areas of the mushroom dissipate heat differently. The changes in mushroom temperature during cooling manifested more irregular thermal gradients when compared to the heating incubation (**Fig S2**). The change in average mushroom temperature as a function of time followed an exponential curve during heating but a linear curve during cooling (**Fig S3**).

**Figure 2.**
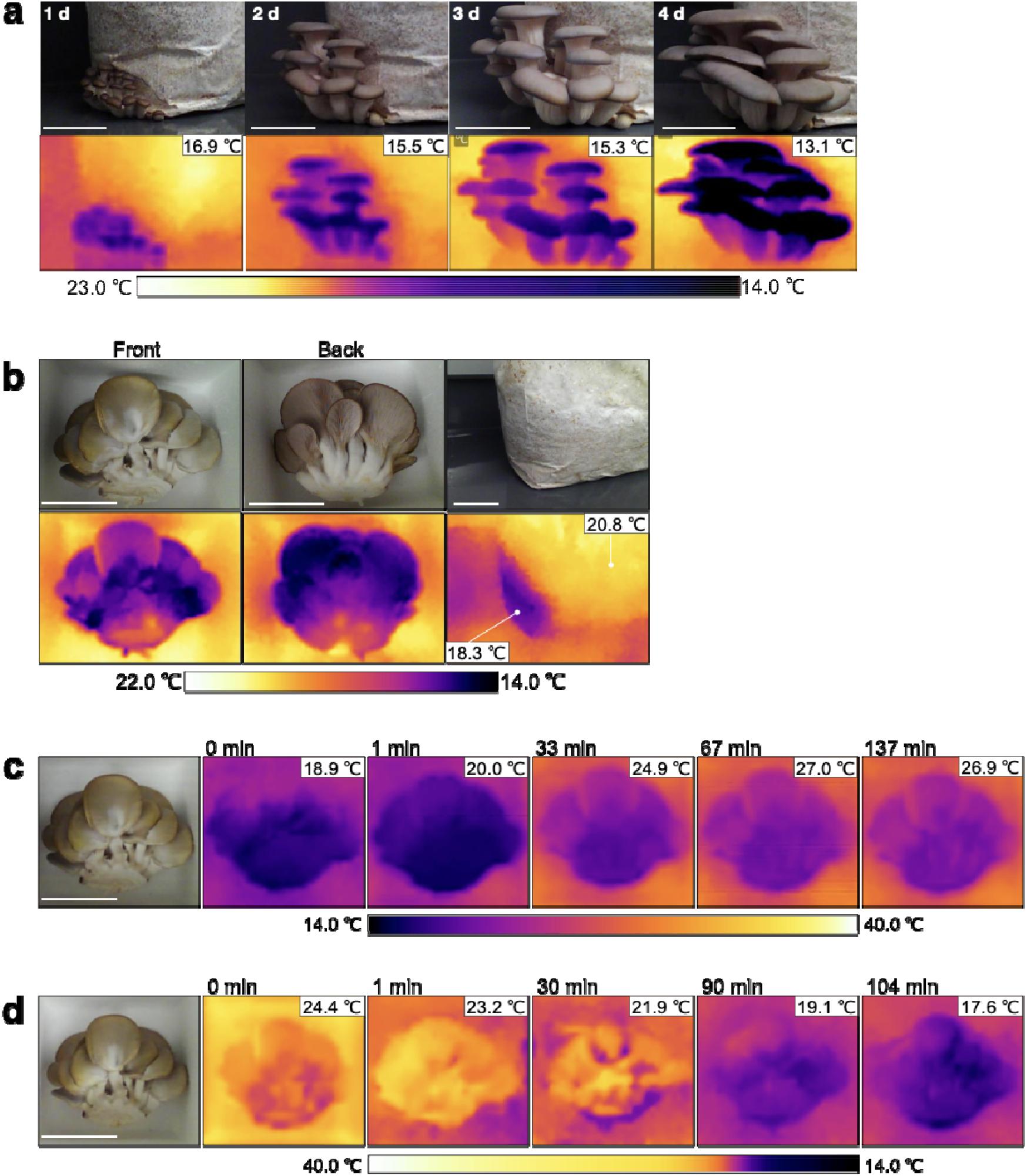
The mushroom temperature during fruiting and during heating and cooling. (**a**) Visible and thermal images of *Pleurotus ostreatus* during fruiting while still attached to its substrate at a temperature-controlled room (22 ± 5°C, 50% RH). Inset temperature values correspond to the lowest temperature signal registered in the thermograph. (**b**) Frontal and back imaging of mushrooms and mycelium bag after detachment on day 4. Thermal imaging of *P. ostreatus* following incubation inside (**c**) warm room at 37 °C, <10% RH followed by incubation inside (**d**) cold room at 4 °C, ~30% RH. Inset temperature values correspond to the lowest and highest temperature signal in the thermographs in (c) and (d), respectively.

### Fungal coldness is mediated via evapotranspiration

Evaporative cooling was confirmed in light and dark substrate-detached *A. bisporus* mushroom pilei by manipulating its water content and ambient temperature-humidity. Dehydrated mushrooms are no longer able to maintain relative colder temperatures, irrespective of ambient temperature (**Fig S4 a-c** and **Table S2)**. We observed similar temperature changes between light and dark mushrooms pilei. The percent mass loss of light and dark *A. bisporus* mushroom pilei following dehydration was 93.6 ± 0.4 % w/w (**Table S3**), demonstrating their high-water content. Dehydration of seven additional wild fungal unidentified specimens also shows high water content ranging from 57 to 92 percent by mass (**Table S3**). Mushrooms warmed slower and reached lower absolute temperatures under a dry environment as compared to a humid environment (**Fig S4 d&e**). Together these results confirmed that mushroom’s relative coldness was mediated by evaporative cooling.

Evaporative cooling in yeast and molds was evident from the condensation of water droplets on the lids above *Cryptococcus neoformans* and *Penicillium* spp. biofilms when grown upright on agar plates (**Fig. 3**). Areas of the agar plate where specimens were not growing show lower levels of condensation. An acapsular mutant of *C. neoformans* showed more and larger water droplets than the encapsulated wildtype strain (**Fig. 3 a&b**). The encapsulated *C. neoformans* colonies are ~90% water, while the acapsular mutant is ~82% (**Table S3**). The mutant strain was also ~1 °C colder than the encapsulated strain. Biofilms of *Penicillium* spp. showed significant condensation of water droplets (**Fig. 3c**), at least ~10 times higher than the surrounding 1.5% agar medium (**Table S4**).

**Figure 3.**
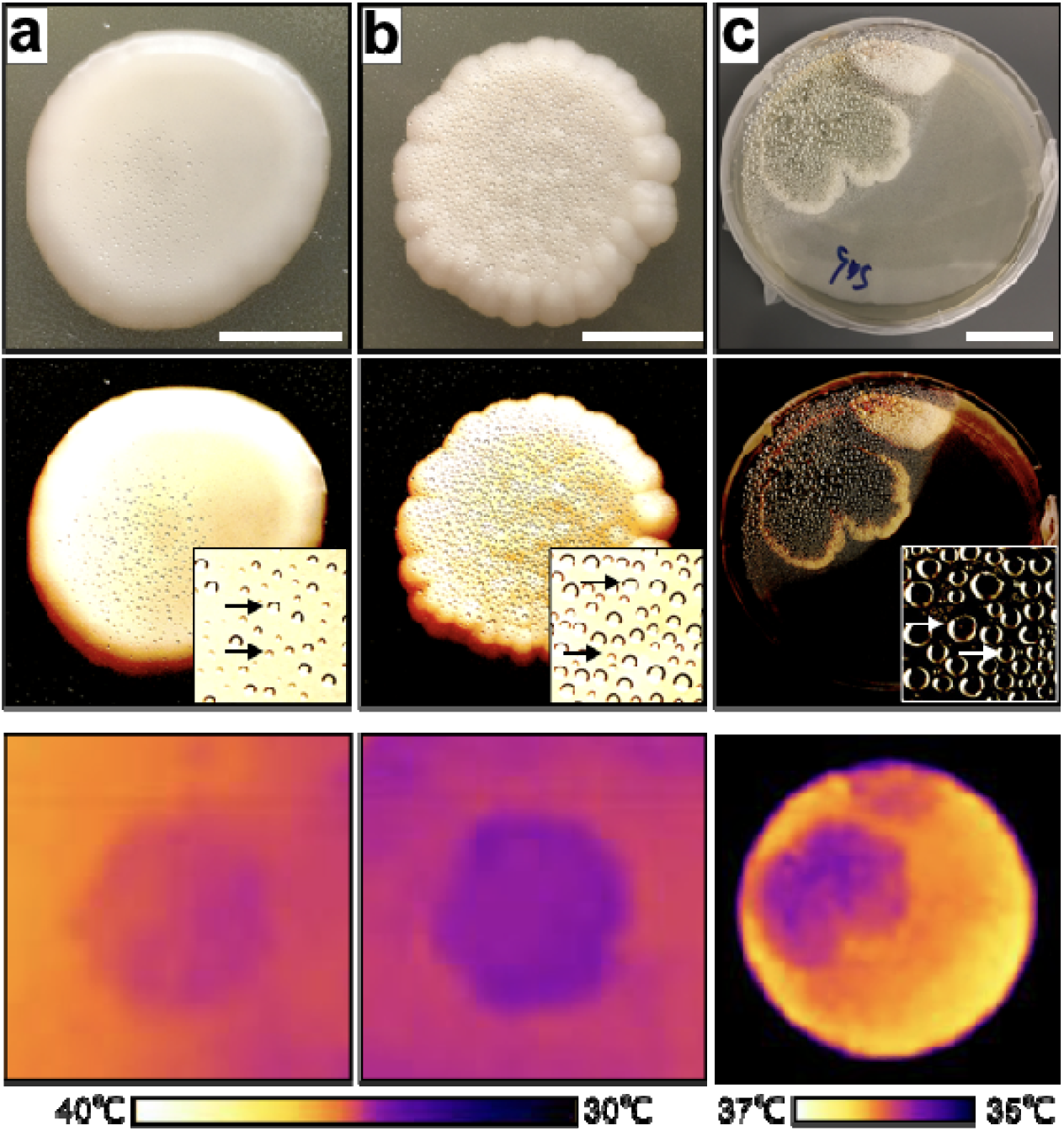
Evaporative cooling in yeast and mold biofilms. Evidence for evaporative cooling is observed from the condensed water droplets at the lid of the petri dish on top of the colony/biofilm. Visible (top and middle) and thermal images (bottom) of (**a**) wildtype H99 *Cryptococcus neoformans;* scale bar 1 cm; (**b**) *cap59* acapsular mutant of *C. neoform*ans; scale bar 1 cm, and (**c**) normal *Penicillium* spp.; scale bar 3 cm. Visible images were altered to increase contrast and help visualize water droplets (middle row).

### A mushroom-based air-cooling device

**Figure 4** shows a diagram of a mushroom-based air-cooling device. We called this prototype device MycoCooler™, which was constructed using a Styrofoam box with a 1-cm diameter inlet aperture and a 2-cm diameter outlet aperture (**Fig S5 a**). An exhaust fan was attached outside the outlet aperture to drive airflow in and out of the box (**Fig. S5 b**). The MycoCooler™ was loaded with ~420 grams of substrate-detached *A. bisporus* mushrooms, closed, and placed inside a larger Styrofoam box (**Fig S5 c**) previously equilibrated inside a warm room (37.8 °C, <10% RH). The temperature inside the closed Styrofoam box decreased from 37.8 °C to 27.8 °C, forty minutes after the addition of mushrooms; cooling approximately 10 °C at ~0.4 °C per min (**Fig 4 b**). In parallel, the humidity increased by ~45% at 1.3 % per min. The box interior reached the coldest temperature at ~60% RH, at which point it started to warm up again to initial temperature values as the humidity continued to increase towards saturation (**Fig 4 b**). From this data, we estimated that 420 grams of *A. bisporus* mushroom pilei have an air-cooling capacity of approximately 20 Watts or 68 BTH/hr. The change in air temperature was proportional to change in humidity, confirming evaporative cooling as the mechanism for mushroom relative coldness. Our MycoCooler™ device provides a proof-of-principle for harnessing mushroom’s cooling capacity for cooling air in enclosed environments.

**Figure 4.**
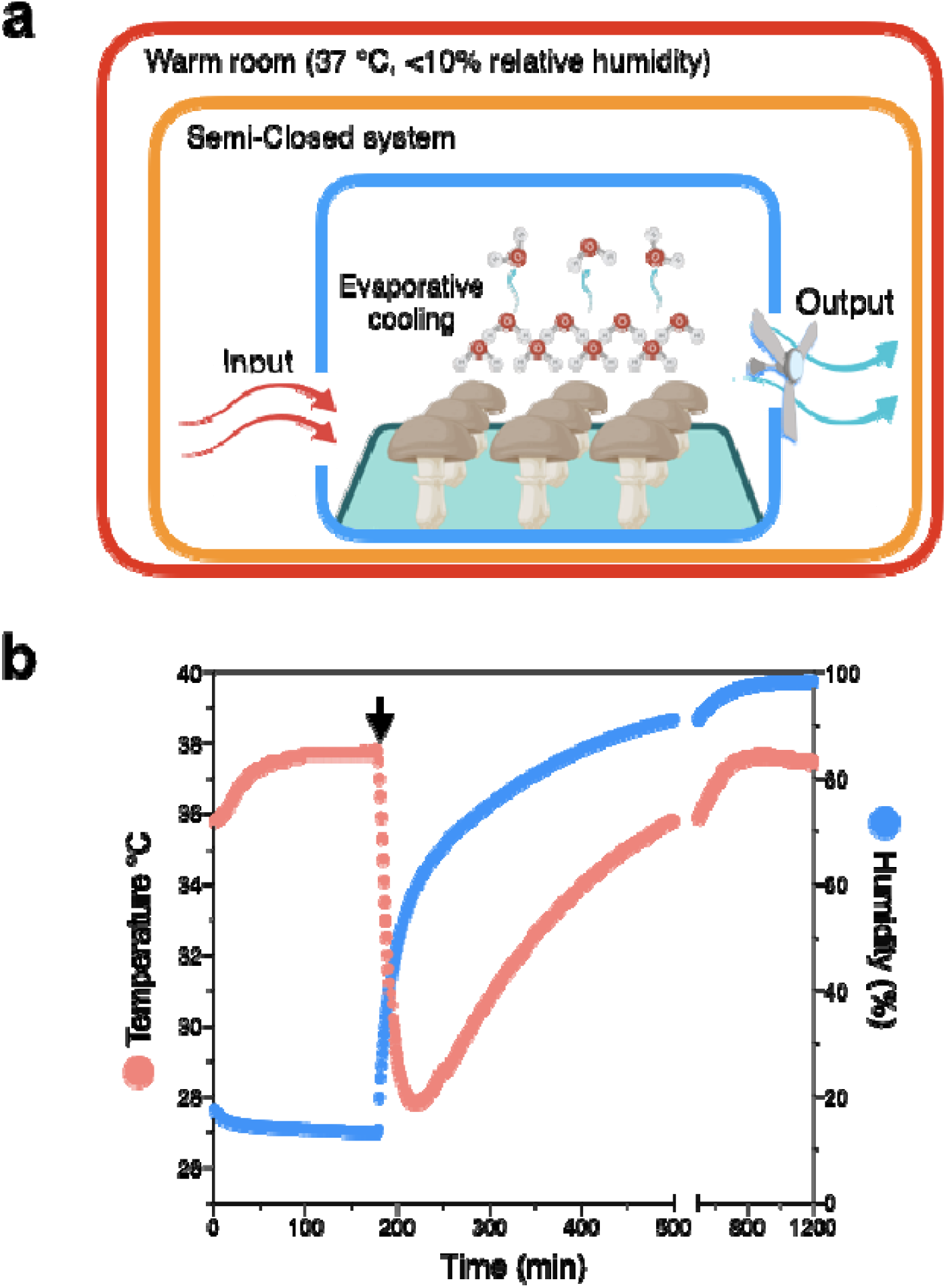
Proof-of-concept of a mushroom-based air conditioning system. (**a**) Prototype diagram model of MycoCooler™ air conditioning system inside a semi-closed system. Warm air enters an insulated chamber containing mushrooms. As the warm air flows inside the chamber, mushroom-mediated evaporative cooling will cool the air. An exhaust fan will push the cooled air through a HEPA filter to limit spore dispersal and enhance air circulation. The fan can be powered via a photovoltaic cell making this system free of carbon emissions. (**b**) Air temperature and relative humidit as a function of time inside a Styrofoam box as the semi-closed system. The black arrow points to the time when commercially available and detached mushrooms were added inside MycoCooler™; once the temperature inside the Styrofoam box semi-closed system reached steady-state.

## Discussion

This study reveals that, in addition to mushrooms, molds and yeasts can also maintain colder temperatures than their surroundings, also via evapotranspiration, suggesting that relative coldness is a general property of the fungal kingdom. This will become clearer as more thermal data about fungal specimens develop in the future. Mushroom coldness occurs throughout the fruiting process, and fruiting areas of mycelium also become colder than non-fruiting areas. We also show that the process of evapotranspiration in yeast and mold biofilms can be measured from the accumulation of water droplets on agar plates. Finally, we provide a proof-of-principle demonstration for a mushroom-based air-conditioning device capable of passively cooling and humidifying the air of a closed environment. The data presented here reveal the cold nature of fungal biology and evaporative cooling as a microbiological mechanism of thermoregulation.

The observations that fungal temperature is colder than the surroundings and correlated with ambient temperature suggest that fungi possess poikilothermic properties. The extent to which fungal temperatures vary with their environmental niche likely involves multiple factors that require further study. The temperature of wild mushrooms relative to ambient temperature varies with the genus, suggesting that there may be species-specific capacities to dissipate heat. The ability to dissipate heat could be related to a variety of factors such as differences in water content, thermal properties (i.e., heat capacity, thermal conductivity), size, color, habitat, lifestyle, and/or phylogenetic relations. These and other factors are important to consider as we learn more about the thermal biology of fungi and how it relates to other kingdoms.

Although fungi tended to be colder than their surroundings, we also observed an instance where a wild dark-colored mushroom appeared warmer than its surroundings. This could be explained by the fact that the mushroom was in its natural open-habitat and exposed to solar radiation, directly or indirectly. By definition, pigments can absorb some fraction of the EM radiation spectrum and lead to temperature increase. Similar to many poikilotherms, fungi also appear to use pigments to increase heat capture from electromagnetic radiation [21,22]. This process of pigment-mediated thermoregulation influences the geographical distribution and thermal adaptation of organisms to changes in climate. This biological phenomenon is discussed as the Gloger’s rule (if also considering precipitation effects) and theory of thermal melanism; all of which have been mainly documented and studied in metazoans [23–25]. With that being said, it is also possible that some fungal organisms can become warmer than their surroundings independently of pigment-mediated heat capture, but we have yet to identify fungal specimens that can maintain relatively warm temperatures even under dark or close-habitat conditions. The identification of heat-producing organisms outside the kingdom of Animalia is rare, but not unprecedented. For example, the identification of heat-producing bacteria [13] suggests that a microbial community can produce enough thermal energy and maintain warmer temperatures than its surroundings.

Mushroom coldness was observed throughout the whole fruiting process. The relative cold temperature of the *P. ostreatus* can be observed since the appearance of the first mushroom pinheads. The observation that the mushroom flush becomes colder as it grows in size, suggests that coldness is related to fungal thermal mass and/or to an unknown age-related structural or physiological process favoring heat loss. The observation that the mushroom is coldest when still attached to the mycelium is consistent with prior observations [18,19] and indicates that heat loss is highest when connected to its substrate via the mycelial network, which permits flow and access to water. This increase in temperature after detachment is also observed in leaves [26].

The observation that the fruiting site of mycelium remained cold after mushroom detachment, suggests that the fungal capacity to cool is also observed at the mycelium level. The cooling of mycelium could be related to mushroom-mediated evapotranspiration or even independently regulated by the mycelium itself. around the fruiting area. Fruiting areas of mycelium could become colder than non-fruiting areas by increasing evapotranspiration rates, even before the appearance of the first fruiting bodies. Further studies are needed to evaluate evapotranspiration during these early stages of the fruiting process and the potential role of mycelium-mediated evapotranspiration in reproductive success and evolution.

The thermal profiles of *P. ostreatus* mushrooms during heating and cooling suggest that heat exchange can vary in discrete areas of the mushroom flush, suggesting that these dissipate heat differently. This could result in areas having different thermal properties, water content, or hyphal structural organizations; all of which could affect evapotranspiration rates. The change in *P. ostreatus* mushroom average temperature as a function of time during heating, followed by cooling incubations, resembles the phenomena of thermal hysteresis, a process where previous heat dissipation events influence subsequent temperature changes. The relatively cold temperatures of hymenium could be explained by the relatively high surface area exposed by lamellae or gills [16]. The high surface structure and organization of hymenium’s lamellae is known to enhance airflow [18,27] and is also expected to enhance heat exchange. Having a relatively cold spore-bearing surface is considered to be important for spore detachment and release [18,19]. Spore discharge is known to be triggered by the mass and momentum transfer of microscopic drops (aka Buller’s drop) on the spore surface [28]. The Buller’s drop is formed by the condensation of moist air in a colder surface [29].

In addition to spore discharge, colder temperatures could also be relevant to fungal sporogenesis. There are multiple examples in nature where sporogenesis is associated with relatively cold temperatures. In mammals, the production of spermatozoa occurs at temperatures colder than the abdominal viscera [30]. In fungi, the frequency of *S. cerevisiae* sporulation and tetrad formation is higher at 22 than at 30 °C [31]. The relation between relatively cold temperatures and sporogenesis suggests that fungal coldness influences reproductive success, by enhancing both the discharge and the production of spores. It also implies that evaporative cooling in fungi is contributing to the thermodynamics regulating spore production and DNA duplication. The relatively cold temperature of mushrooms may also be attractive to insects which also contribute to spore dispersal. Considering that thermoregulation is a general defense strategy to tolerate infectious diseases [32], the relatively cold temperatures of fungi should be considered to understand infection by bacteria or mycoviruses. The ability to dissipate heat via evapotranspiration may also be important for fungal temperature priming and memory [33]. In this case, evapotranspiration may provide a window of opportunity for the fungus to adapt to different levels of temperature stress, where a priming temperature stress renders the fungus to be better prepared for more severe temperature stresses. The ability of fungal organisms to tolerate higher temperatures represents a serious public health concern considering that new pathogenic fungal species could emerge as a result of climate change [34].

Our indoor measurements of lightly and darkly pigmented *A. bisporus* detached mushroom pilei confirmed that evaporative cooling accounts for their relatively cold temperatures. Both light and dark pilei exhibited similar temperatures changes suggesting that pigmentation may not have an effect on heat dissipation under such conditions or that the effect is too close to our limits of thermal detection. The high-water content of mushrooms is consistent with previous reports [35] and the high water content of yeasts also implies that fungal thermal properties must be largely determined by water. The evapotranspiration capacity of mushrooms was reported to be higher than in some plants [17,19]. In plants, evapotranspiration occurs mainly at the leaves, and the rate of evapotranspiration is regulated at the level of stomata. An analogous cellular structure dedicated to regulating evapotranspiration in mushrooms has not been identified. It would be interesting to compare the temperature and evapotranspiration capacity of mushrooms with other plants and fruits that are also highly hydrated (e.g. aloe vera, ~98% water). There must be something about the structure and composition of mushrooms that accounts for their relatively high capacity of evapotranspiration. Evapotranspiration in fungi provides a passive physiological mechanism for dissipating heat and although it appears to be dominant, it may not be the only biological mechanism contributing to fungal coldness.

Evapotranspiration in yeast and molds was evident from the water condensing above biofilms, which maintained colder temperatures relative to the surrounding agar, which is also mostly water. It is important to note that the relatively cold temperature of colonies and biofilms were only detectable following long or a transient incubation of plates in a relatively warm and dry environment. When at steady states in ambient temperature, we were not able to detect a clear temperature difference between the colony or biofilm versus the surrounding agar with the current contrasting thermal resolution. The use of microbolometers with higher resolving power will confirm the temperature difference between fungal colonies and biofilms relative to the surrounding agar or substrate at transient versus steady states.

We think that water condensation above fungal colonies and biofilms could be used as a parameter to evaluate evapotranspiration, water flow, or metabolism. An index of evapotranspiration derived from water condensation may be used to screen for genetic determinants of thermoregulation in yeast. Our data suggest that a gene deletion can result in notable differences in both evapotranspiration and temperature. Our data show that the acapsular mutant of *C. neoformans* contained ~10% less water mass but exhibited colder temperatures and more evapotranspiration relative to the normal encapsulated strain. The water difference between the encapsulated and acapsular mutant biofilms could also be explained by the presence of the *C. neoformans* polysaccharide capsule, which is mostly water [19]. Hydrated polysaccharide capsules are considered to prevent desiccation in environmental microorganisms by retaining water [20]. Such water retention may be limiting the rate of evapotranspiration in *C. neoformans*.

Our data shows that mushrooms can be used for air cooling and conditioning, consistent with previous reports [18]. The relatively high transpiration rate of mushrooms could be exploited to develop a natural and passive air-conditioning system. The Mycocooler™ prototype presented here produced a cooling capacity of 1002.3 BTU/hr. For reference, a 100-150 ft^2^ room often requires a cooling unit with a capacity of 5,000 BTUs/hr [36]. Even though these kinds of traditional window air conditioners present at least five times more cooling capacity, these units also weight >20 times more. This suggests that a mushroom-based air condition device could provide superior cooling capacity per unit weight, although other features like performance, volume, or aroma may also be potential tradeoffs when comparing it to traditional air conditioning units. Mushroom-based air cooling depends on relative humidity, and for detached *A. bisporus* mushroom pilei, cooling is compromised at a relative humidity close to 60%. Better results could be achieved using mushroom species with higher transpiration rates, still attached to their mycelium, and on a design that controls the accumulation of air moisture. Mushrooms can be used not only to cool the surrounding air but also to humidify and even purify with low electrical energy consumption and/or CO_2_ emissions. These findings suggest the possibility of using massive myco-cultures to reduce the temperatures of locales and help mitigate local global warming trends.

Given that fungi live on soils and comprise 2% of the Earth’s biomass [4], that fungi are 2-5 °C cooler than their environment, that the average surface temperature of the Earth is ~15 °C [37], and assuming linearity in heating and cooling, we used two methods for roughly estimating what the temperature of the planet would be without fungi. One method determined that the Earth would be 0.04-0.1 °C warmer, while the second method determined that the Earth would be 0.44-0.58 °C warmer. It is important to note that these calculations are also based on the assumption that most of the total fungal mass is composed of fruiting bodies or structures that exhibit similar transpiration rates as the fungal species documented here. The cooling activity of fungal mycelium and how it compares to other fungal life forms remains to be determined. These methods both present interesting postulates to evaluate the impact of fungi on global temperatures, but their disparities also confirm their approximate nature.

The ability of certain fungi to trap carbon in soil has already pointed to their potential role in mitigating climate change [38]. The identification of fungi as potential heat sinks raises additional support for their role in global and local temperature regulation. Within the realm of ecology, fungal-mediated cooling raises several questions about their effects on a microclimate, as a symbiont, and in an ecosystem. One question that could confirm this study’s findings on a larger scale would be whether ecosystems with higher fungal biomass exhibit lower ambient temperatures, as seen in areas with large amounts of plant life or water bodies, for example.

In conclusion, this study reveals the cold nature of fungi and evaporative cooling as a fundamental mechanism for thermoregulation for this kingdom. The relative cold temperature of fungi (or fungal hypothermia), implies that their heat loss is much greater than the production of heat via metabolism. Their relative cold temperatures also imply that the flow of surrounding thermal energy will move towards the fungus, turning it into some sort of heat sink. The high-water content and evapotranspiration rate of fungi implies that their molecular composition and structure enables the efficient transfer of thermal energy and water. Infrared imaging provides a powerful tool to study the thermal biology of mushrooms, mycelium, molds, yeasts at the community level. A yeast model system to study thermal biology could allow the screening of genetic and epigenetic mechanisms regulating thermodynamics and thermal fitness. Understanding how fungal organisms dissipate heat is relevant to a world of applications, from novel biotechnologies to medical innovations, and sustainable energy **Figure 5**.

**Figure 5:**
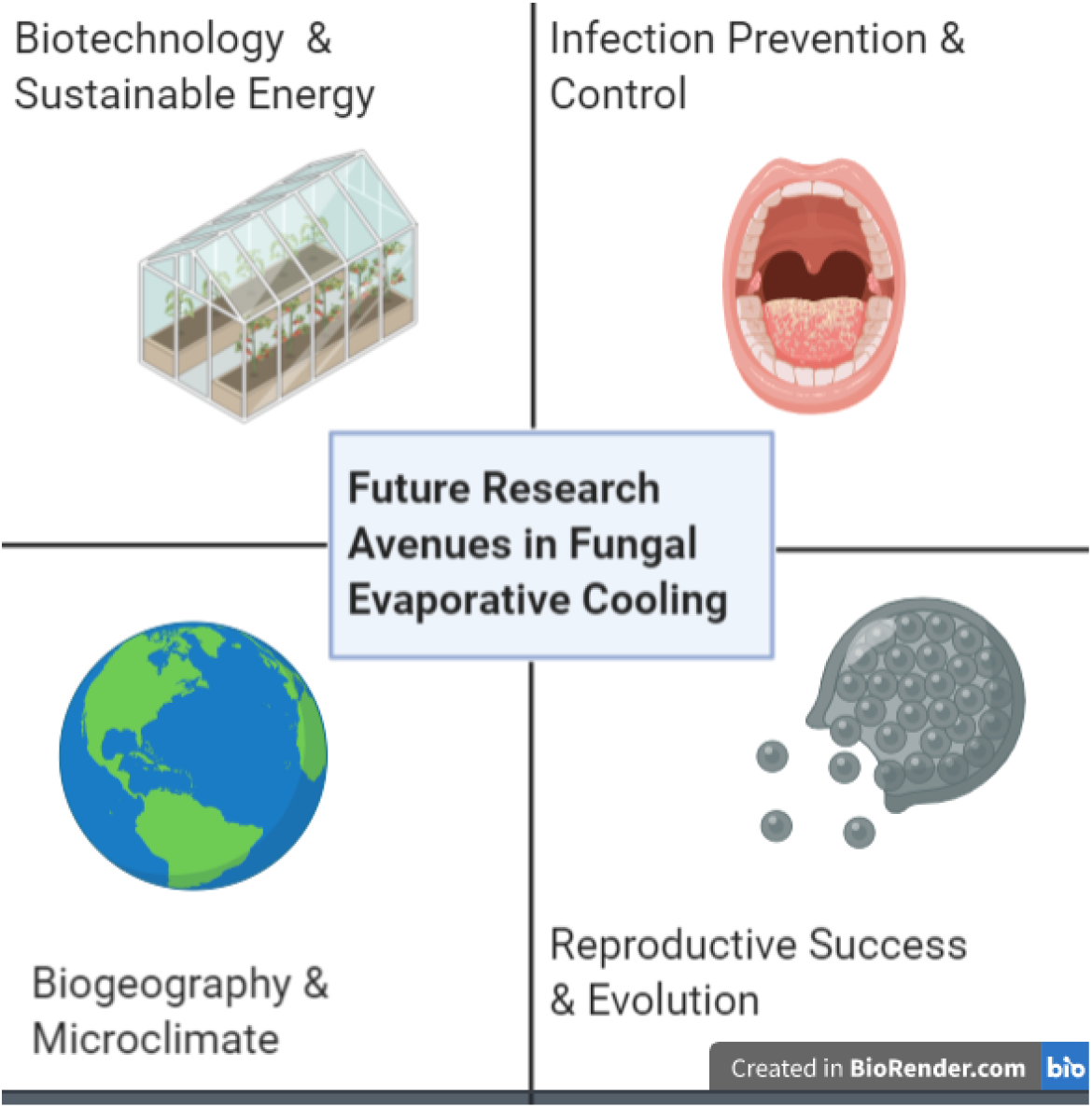
Areas of future research built on the theory of fungal evaporative cooling. Each sector represents a different area of application and further questions with supporting graphics. Pictures are as follows: Infection Prevention & Control (oral candidiasis infection), Reproductive Succes & Evolution (coccidioides releasing spores), Biogeography & Microclimate (Earth), and Biotechnology & Sustainable Energy (greenhouse).

## Methods

All wild mushrooms specimens were found in Lake Roland Park in the state of Maryland during the evenings of July 5, 6, and 9^th^ of 2019. Partial and non-official identification of specimens was made based on a visual inspection and photograph analysis via crowdsourcing the Internet. *Candida* spp., *Clodosporium* spp., and *Penicillium* spp. were obtained from mosquito gut isolates by the Dimopoulos Laboratory at the MMI Dept. *Rhodotorula mucilaginosa* was isolated from a contaminated YPD agar plate in our laboratory. *C. neoformans* Serotype A strain H99 (ATCC 208821), acapsular cap59 mutant yeasts, and *Penicillium* spp. molds were grown on Sabouraud Dextrose agar or liquid media for 3-5 days at 30 °C and 24 °C, respectively. *Pleurotus ostreatus* was purchased from The Mushroomworks (Baltimore, MD) as an already-inoculated substrate contained in a 6-pound clear filter patch bag. Fruiting was triggered by making a single 4-inch side cut on the bag and letting it standstill at 24 °C for seven days. Mushroom flush was detached from mycelium on day four after it started fruiting. Light and dark *Agaricus bisporus* were purchased from New Moon Mushrooms (Mother Earth, LLC., Landenberg, PA, USA) and L. Pizzini & Son, Inc. (Landenberg, PA, USA), respectively.

### Thermography

Wild mushroom temperatures were measured in their natural habitats while still attached to their natural substrates using a FLIR C2 IR camera (FLIR Systems, Wilsonville, OR). The camera specifications are 80×60-pixel thermal resolution; 640×480-pixel visual camera resolution; 7.5-14 μm spectral range of camera detector; object temperature range of −10 to 150 °C, accuracy ±2 °C or 2%, whichever is greater, at 25 °C nominal; thermal sensitivity: <0.10 °C; adjusted emissivity to 0.96. The ambient temperature was derived from a card-containing black vinyl electrical tape with an emissivity of 0.96 and aluminum foil (emissivity 0.03). The black tape and aluminum foil were included in the picture as a reference for ambient and reflective temperature readings, respectively (**Fig. S6 a**).

The efficacy of the black vinyl tape in reproducing ambient temperatures was tested by thermal imaging of the reference card following ~20 minutes incubation inside three temperature-controlled rooms, set to approximately 5, 25, and 38 °C. The temperature readings obtained from the black tape using the thermal camera were 5.2 (5.2/5.4 min/max), 25.5 (25.5/25.5 min/max), and 37.9 (37.7/38.1 min/max) °C, respectively; demonstrating that black tape radiative temperature corresponded to the ambient temperature. The temperature readings between the thermal camera and a mercury thermometer matched clearly (**Fig. S6 b**), confirming that black tape radiative temperature matched the temperature of rooms, hence serving as a useful reference for ambient temperature.

The thermography of yeast and molds colonies/biofilms was done similar to was described previously [21]. Thermal images of yeast, molds, and commercial mushrooms (*P. ostreatus* and *A. bisporus*) were taken inside a white Styrofoam box (30 × 27 × 30 mm, and 3.5 mm wall thickness) to prevent heat loss and radiation noise from surroundings. Before imaging yeasts and mold specimens, the sample plates were incubated at 45 or 37 °C. This incubation was required to detect a temperature difference between the colony and the agar as dictated by our thermographic resolution. Following a 1-hour incubation period, the yeast/mold containing plates were immediately transferred inside a Styrofoam box. Next, the box was closed with a lid having a hole fitted to a FLIR C2 IR camera (FLIR Systems, Wilsonville, OR). The camera detector is set at 2.5 nm distance from the specimen. The temperature of the substrate-detached *P. ostreatus* mushroom flush was monitored during heating and cooling by placing the mushroom inside a warm room (37 °C, <10% RH) or cold room (4 °C, ~30% RH) for 137 and 104 minutes, respectively. Thermal images of mushroom flush were taken inside the Styrofoam box at different time intervals. All apparent temperatures of yeasts, molds, and mushrooms were obtained from infrared images using the FLIR Tool analysis software Version 5.13.17214.2001. Plot profiles of thermal images were obtained using the ImageJ software.

### Water condensation of fungal biofilms

*C. neoformans* yeast biofilms were prepared by spotting 25 μL of a liquid 2-day old pre-culture on Sabouroaud agar medium. The liquid pre-cultures were inoculated from frozen stock and grown for two days at 30 °C (shaking at 180 rpm). *Penicillium* spp. biofilm was naturally formed by inoculating on a Sabouroaud agar plate. Yeast and mold-inoculated plates were grown upright at 24 °C for 1-2 weeks or until water condensation on the lids became visible. The amount of condensed water at the lid above a mold’s biofilm or plain agar was collected using a Steriflip® filter vacuum unit (Millipore Sigma) connected to two 50 mL conical tubes; one at each end. Suction was achieved by connecting a small tubing across the filter into one of the conical tubes. A pipet tip connected at the end of tubing facilitated the aspiration of condensed water droplets on the lid and its collection into one of the 50 mL conical for weighting. The water mass was normalized by the condensation area on the lid, which was estimated from digital images using the ImageJ software.

### Mushroom dehydration

Mushrooms were dehydrated for five days using a freeze-drying system (Labcono, Kansas City, MO).

### Thermocouple-thermometry of pilei

To monitor the temperature of *A. bisporus* mushrooms as a function of time, mushroom pilei of equal masses, kept at 24 °C, were placed on glass trays inside ziplock clear bags (one mushroom per bag). One bag contained 40 grams of desiccating-anhydrous indicating Drierite (W.A. Hammond Drierite Company, LTD), and a second contained 40 grams of distilled water. Thermocouple detectors (K-type) were submerged inside each mushroom cap (centered from top). Bags were then closed and placed inside a warm room (37 °C, <10 % RH), and temperature readings were recorded every second using the Amprobe TMD-56 thermometer (0.05% accuracy) connected to a computer. Each mushroom sample was measured individually inside the warm room.

### A mushroom-based cooling device

A prototype for a mushroom-based air-cooling device or MycoCooler™ is shown in (**Figure 4 a**). The prototype device was made using a Styrofoam box with dimensions 20 × 21 × 21 cm or a total volume of 8820 cm^3^. An inlet aperture of 1-cm diameter and an outlet aperture of 2-cm diameter at opposite ends of the box allowed airflow inside and outside the box containing mushrooms (**Fig S5 a**). An exhaust fan (Noctua NF-P12) was glued outside the box on top of the outlet aperture to facilitate the circulation of air (air flow rate of approximately in and out the MycoCooler™ (**Fig S5 b**). Approximately, 420 grams of fresh and substrate-detached *A. bisporus* mushrooms were placed inside the MycoCooler™ box, which was then closed, and placed inside a larger Styrofoam box with dimensions (30.48 × 30.48 × 30.48 cm or a total volume of 28.32 L (28,316.85 cm^3^). This larger box was maintained inside a warm room (37 °C, <10 % RH) throughout the experiment. The MycoCooler™ containing mushrooms was enclosed inside the larger box once the temperature and humidity values reached a steady state. The temperature and relative humidity inside the larger Styrofoam box (**Figure S5 a**) were recorded every minute using an Elitech GSP-6 data logger has a temperature accuracy of ±0.5 °C (−20~40 °C) and humidity range 10%~90% and an accuracy of ±3% RH (25 °C, 20%~90% RH).

### Quantification and statistical analysis

Details for each statistical analysis, precision measures, the exact value of n (and what n represents; sample size and the number of replicates) for all shown data can be found in the figure legends. We used an alpha level of 0.05 for all statistical tests.

To calculate the mushroom’s cooling capacity, we used the MycoCooler™ temperature change data to estimate the cooling capacity of 420 g of *A. biosporus* mushroom pilei. Cooling capacity was calculated using the energy equation for heat transfer Q=mC_p_ΔT, where *m* is the mass flow rate of the air in kg/s, *C_p_* is the specific heat capacity of air in kJ/kg*K, and Δ*T* is the temperature difference in Kelvin. The mass flow rate was obtained by multiplying the density of air at 37 °C and 1 atm (1.138 kg/m^3^) by the fan flow rate taken from equipment specifications (0.0256 m^3^/s). This results in a mass flow rate of 0.0292 kg/s, that if multiplied by the heat capacity nominal value of air at 37 °C (1.006 kJ/kg*K) and the absolute air temperature difference of the enclosed system before and 45 minutes after the addition of mushrooms (27.8 °C+273.15 K)-(37.8 °C+273.15 K)=10.0 K. This yields a heat transfer or cooling capacity of ~ 293.8 Watts or 1,002.3 British Thermal Units per hour (BTU/h). For reference, a 100-150 ft^2^ room often needs a cooling capacity of about 5,000 BTUs/hr [36]. The cooling capacity divided by the mass of the mushrooms 0.42 kg yields 699.4 Watts/kg or 1082.5 BTU/hr/lb. When taking the mass of the MycocoolerTM and mushrooms (0.71 kg) into consideration, the values become 412.2 Watts/kg or 638.0 BTU/hr/lb. In contrast, a brief survey of popular 5000 BTU air conditioners determined that these units weigh about 18 kg, yielding a BTU/hr/lb of 125. Although the MycoCoolerTM produces an impressive cooling capacity per unit weight (>20 times more), the relative volume may be a potential tradeoff in comparing it to traditional air conditioning units.

To estimate the temperature of the Earth without fungi, we used two different calculation methods. The first considered the average temperature difference of wild mushrooms, which ranged from ~2-5 °C, and multiplied that by the estimated proportion of fungal biomass ~2% [4], such that the temperature associated with fungi would be ~0.04 to 0.1 °C. We then estimated the global mean surface temperature without the fungal biomass, X=15 °C+(0.04 or 0.1 °C), such that global temperatures would be ~15.04°C or ~15.1°C, or ~0.3-0.7% warmer. This calculation is based on the assumption that all fungi are not in the substrate, have similar metabolic properties as mycelia, and are heat sinks due to their evaporative cooling properties.

The second estimation method used considered the immense amount of water within fungal organisms, and how the atmosphere may warm in the absence of that water. We began by considering the total mass of water contained in fungal bodies. As fungi make up about 12 Gt global carbon, are approximately 14.29% Carbon by wet weight [39], and ~75% water by weight, approximately 6.3×10^19^ kg of water is contained in fungal bodies. This quantity was then multiplied by the specific heat of water (4.18 J/kg*°C) and the upper and lower estimates for average fungal temperature (average global temperature 15°C-[2,5]) to yield a range of heat storage between 2.6×10^21^ - 3.4×10^21^ Joules. By multiplying the specific heat of air at 300 K by the estimated mass of the atmosphere [40], we obtained an estimated heat capacity value for the atmosphere of 5.961×10^21^ J/°C. In dividing the energy storage range by this value, we determined that the global increase in temperature due to an absence of fungi could range from 0.44 °C to 0.58 °C.

## Acknowledgments and Funding

The research was supported by the National Institutes of Health (R01 AI052733). We thank Teporah Bilezikian for helping with fieldwork and the identification of wild mushroom species. We also thank members of The Baltimore Fungal Group and The Casadevall Laboratory, particularly to Samuel dos Santos, Daniel Smith and Zach Stolp, for valuable input.

## Bibliography

1. Gates, D. M. 1980 Biophysical Ecology. New York, NY: Springer New York.(doi:10.1007/978-1-4612-6024-0)

2. Lineweaver, C. H. & Egan, C. A. 2008 Life, gravity and the second law of thermodynamics. Phys. Life Rev. 5, 225–242. (doi:10.1016/j.plrev.2008.08.002)

3. Clarke, A. 2017 Principles of thermal ecology: temperature, energy, and life. Oxford University Press.(doi:10.1093/oso/9780199551668.001.0001)

4. Bar-On, Y. M., Phillips, R. & Milo, R. 2018 The biomass distribution on Earth. Proc. Natl. Acad. Sci. USA 115, 6506–6511. (doi:10.1073/pnas.1711842115)

5. Mahan, J. R. & Upchurch, D. R. 1988 Maintenance of constant leaf temperature by plants—Hypothesis-limited homeothermy. Environ Exp Bot 28, 351–357. (doi:10.1016/0098-8472(88)90059-7)

6. McNaughton, S. J. 1972 Enzymic thermal adaptations: the evolution of homeostasis in plants. Am. Nat. 106, 165–172. (doi:10.1086/282759)

7. Helliker, B. R. & Richter, S. L. 2008 Subtropical to boreal convergence of tree-leaf temperatures. Nature 454, 511–514. (doi:10.1038/nature07031)

8. Wang, K. & Dickinson, R. E. 2012 A review of global terrestrial evapotranspiration: Observation, modeling, climatology, and climatic variability. Rev. Geophys. 50. (doi:10.1029/2011RG000373)

9. Jackson, R. D. 1982 Canopy temperature and crop water stress. pp. 43–85. Elsevier.(doi:10.1016/B978-0-12-024301-3.50009-5)

10. Doughty, C. E. & Goulden, M. L. 2008 Seasonal patterns of tropical forest leaf area index and CO_2_ exchange. J. Geophys. Res. 113. (doi:10.1029/2007JG000590)

11. Still, C., Powell, R., Aubrecht, D., Kim, Y., Helliker, B., Roberts, D., Richardson, A. D. & Goulden, M. 2019 Thermal imaging in plant and ecosystem ecology: applications and challenges. Ecosphere 10. (doi:10.1002/ecs2.2768)

12. Dusenbery, D. B. 2011 Living At Micro Scale: The Unexpected Physics Of Being Small. Harvard University Press.

13. Tabata, K., Hida, F., Kiriyama, T., Ishizaki, N., Kamachi, T. & Okura, I. 2013 Measurement of soil bacterial colony temperatures and isolation of a high heat-producing bacterium. BMC Microbiol. 13, 56. (doi:10.1186/1471-2180-13-56)

14. Meyer, V. et al. 2020 Growing a circular economy with fungal biotechnology: a white paper. Fungal Biol. Biotechnol. 7, 5. (doi:10.1186/s40694-020-00095-z)

15. Fisher, M. C. et al. 2020 Threats posed by the fungal kingdom to humans, wildlife, and agriculture. MBio 11. (doi:10.1128/mBio.00449-20)

16. Fischer, M. W. F. & Money, N. P. 2010 Why mushrooms form gills: efficiency of the lamellate morphology. Fungal Biol 114, 57–63. (doi:10.1016/j.mycres.2009.10.006)

17. Mahajan, P. V., Oliveira, F. A. R. & Macedo, I. 2008 Effect of temperature and humidity on the transpiration rate of the whole mushrooms. J. Food Eng. 84, 281–288. (doi:10.1016/j.jfoodeng.2007.05.021)

18. Dressaire, E., Yamada, L., Song, B. & Roper, M. 2016 Mushrooms use convectively created airflows to disperse their spores. Proc. Natl. Acad. Sci. USA 113, 2833–2838. (doi:10.1073/pnas.1509612113)

19. Husher, J., Cesarov, S., Davis, C. M., Fletcher, T. S., Mbuthia, K., Richey, L., Sparks, R., Turpin, L. A. & Money, N. P. 1999 Evaporative cooling of mushrooms. Mycologia 91, 351–352. (doi:10.1080/00275514.1999.12061025)

20. Paul Stamets 2016 Thermal Amanita muscaria.

21. Cordero, R. J. B., Robert, V., Cardinali, G., Arinze, E. S., Thon, S. M. & Casadevall, A. 2018 Impact of yeast pigmentation on heat capture and latitudinal distribution. Curr. Biol. 28, 2657–2664.e3. (doi:10.1016/j.cub.2018.06.034)

22. Krah, F.-S. et al. 2019 European mushroom assemblages are darker in cold climates. Nat. Commun. 10, 2890. (doi:10.1038/s41467-019-10767-z)

23. Clusella Trullas, S., van Wyk, J. H. & Spotila, J. R. 2007 Thermal melanism in ectotherms. J Therm Biol 32, 235–245. (doi:10.1016/j.jtherbio.2007.01.013)

24. Stuart-Fox, D., Newton, E. & Clusella-Trullas, S. 2017 Thermal consequences of colour and near-infrared reflectance. Philos. Trans. R. Soc. Lond. B, Biol. Sci. 372. (doi:10.1098/rstb.2016.0345)

25. Delhey, K. 2017 Gloger’s rule. Curr. Biol. 27, R689–R691. (doi:10.1016/j.cub.2017.04.031)

26. Utsumi, Y. et al. 2019 Acetic Acid Treatment Enhances Drought Avoidance in Cassava (Manihot esculenta Crantz). Front. Plant Sci. 10, 521. (doi:10.3389/fpls.2019.00521)

27. Deering, R., Dong, F., Rambo, D. & Money, N. P. 2001 Airflow patterns around mushrooms and their relationship to spore dispersal. Mycologia 93, 732–736. (doi:10.1080/00275514.2001.12063204)

28. Turner, J. C. R. & Webster, J. 1991 Mass and momentum transfer on the small scale: how do mushrooms shed their spores? Chem Eng Sci 46, 1145–1149. (doi:10.1016/0009-2509(91)85107-9)

29. Webster, J., Davey, R. A. & Ingold, C. T. 1984 Origin of the liquid in Buller’s drop. Transactions of the British Mycological Society 83, 524–527. (doi:10.1016/S0007-1536(84)80055-8)

30. Glover, T. D. & Young, D. H. 1963 Temperature and the production of spermatozoa. Fertil. Steril. 14, 441–450. (doi:10.1016/s0015-0282(16)34929-9)

31. Codón, A. C., Gasent-Ramírez, J. M. & Benítez, T. 1995 Factors which affect the frequency of sporulation and tetrad formation in Saccharomyces cerevisiae baker’s yeasts. Appl. Environ. Microbiol. 61, 630–638. (doi:10.1128/AEM.61.2.630-638.1995)

32. Schieber, A. M. P. & Ayres, J. S. 2016 Thermoregulation as a disease tolerance defense strategy. Pathog Dis 74. (doi:10.1093/femspd/ftw106)

33. Andrade-Linares, D. R., Veresoglou, S. D. & Rillig, M. C. 2016 Temperature priming and memory in soil filamentous fungi. Fungal Ecology 21, 10–15. (doi:10.1016/j.funeco.2016.02.002)

34. Casadevall, A., Kontoyiannis, D. P. & Robert, V. 2019 On the Emergence of Candida auris: Climate Change, Azoles, Swamps, and Birds. MBio 10. (doi:10.1128/mBio.01397-19)

35. Vaz, J. A., Barros, L., Martins, A., Santos-Buelga, C., Vasconcelos, M. H. & Ferreira, I. C. F. R. 2011 Chemical composition of wild edible mushrooms and antioxidant properties of their water soluble polysaccharidic and ethanolic fractions. Food Chem. 126, 610–616. (doi:10.1016/j.foodchem.2010.11.063)

36. EnergyStar.gov In press. Room Air Conditioner. Buying Guidance.

37. Hansen, J., Ruedy, R., Sato, M. & Lo, K. 2010 GLOBAL SURFACE TEMPERATURE CHANGE. Rev. Geophys. 48. (doi:10.1029/2010RG000345)

38. Averill, C., Turner, B. L. & Finzi, A. C. 2014 Mycorrhiza-mediated competition between plants and decomposers drives soil carbon storage. Nature 505, 543–545. (doi:10.1038/nature12901)

39. Fidanza, M. A., Sanford, D. L., Beyer, D. M. & Aurentz, D. J. 2010 Analysis of fresh mushroom compost. Horttechnology 20, 449–453. (doi:10.21273/HORTTECH.20.2.449)

40. Trenberth, K. E. & Smith, L. 2005 The mass of the atmosphere: A constraint on global analyses. J. Clim. 18, 864–875. (doi:10.1175/JCLI-3299.1)

