## Supplemental Information for "Fungi are colder than their surroundings"

**Supporting Information**

This is the supplemental material for “Fungi are colder than their surroundings” that includes 3 tables and 7 figures.

**Table S1.** Apparent temperature of two yeast and two molds colonies/biofilms grown in agar plates, and 21 wild mushrooms pilei of fungal specimens in their natural environment. For yeast and mold specimens, the surrounding temperature corresponds to the agar temperature derived from infrared images. For wild mush mushroom specimens, the surrounding temperature corresponds to the ambient air temperature. The stalk temperature (in parenthesis) was also recorded for some mushroom specimens.

| **Fungal Specimen** | **ID#** | **fungal specimen °C** | | | **Surrounding temperature °C** | | | **Mean Difference** (absolute) |
| --- | --- | --- | --- | --- | --- | --- | --- | --- |
|  |  | average | max | min | average | max | min |  |
| *Yeast and molds* | | | | | | | | |
| *Candida* spp. | 1 | 36.9 | 37.4 | 36.6 | 38.2 | 38.15 | 38.1 | 1.3 |
| *Candida* spp. | 2 | 40.6 | 40.7 | 40.6 | 41.3 | 41.9 | 40.7 | 0.7 |
| *Cladosporium sphaerospermum* | 3 | 38.7 | 39.4 | 38.2 | 41.3 | 41.9 | 40.7 | 2.6 |
| *Penicillium* spp. | 4 | 28.7 | 29 | 28.2 | 30.6 | 30.7 | 30.4 | 1.9 |
| *Rhodotorula mucilaginosa* | 5 | 33.8 | 33.8 | 33.7 | 36.9 | 37 | 36.9 | 3.1 |
| *Mushroom caps (and stalks)* | | | | | | | | |
| *Amanita* spp. | 6 | 23.8 (23.9) | 24.1 (23.8) | 23.7 (24.1) | 26.1 | 26.2 | 26.1 | 2.3 |
| *Pleurotus ostreatus* | 7 | 26.6 | 26.7 | 26.6 | 31.7 | 31.9 | 31.3 | 5.1 |
| *Amanita muscaria* | 8 | 24.6 (24.4) | 24.4 (24.1) | 24.7 (24.6) | 27.3 | 27.2 | 27.4 | 2.7 |
| *Amanita brunnescens* | 9 | 24.7 (24.9) | 24.7 (24.8) | 24.8 (25.0) | 26.5 | 26.4 | 26.6 | 1.8 |
| *Russula* spp. | 10 | 24.8 (24.9) | 24.7 (24.8) | 25.2 (25.1) | 27.4 | 27 | 27.6 | 2.6 |
| *Boletus separans* | 11 | 26.2 | 25.9 | 26.5 | 29.5 | 29.1 | 30 | 3.3 |
| *Russula* spp. | 12 | 24.1 (24.5) | 24.0 (24.4) | 24.3 (24.5) | 26.7 | 26.5 | 26.8 | 2.6 |
| *Amanita* spp. | 13 | 24.6 (24.5) | 24.5 (24.3) | 24.8 (24.7) | 26.3 | 26.2 | 26.6 | 1.7 |
| *Thelephora* spp. | 14 | 25.6 | 25.3 | 25.9 | 28.7 | 28.3 | 28.9 | 3.1 |
| *Cerrena unicolor* | 15 | 29 | 28.6 | 29.2 | 34.1 | 33.2 | 34.6 | 5.1 |
| *Cantharellus* spp. | 16 | 23.8 | 23.7 | 23.9 | 25.8 | 25.7 | 26 | 2 |
| *Russula* spp. | 17 | 24.4 (24.2) | 24.2 (24.1) | 24.7 (24.2) | 25.8 | 25.7 | 25.9 | 1.4 |
| *Hortiboletus* spp. | 18 | 24.2 (24.0) | 24.1 (24.0) | 24.3 (24.0) | 26.7 | 26.5 | 26.8 | 2.5 |
| *Marasmius capillaris* | 19 | 24.5 | 24.4 | 24.6 | 26.4 | 26.3 | 26.5 | 1.9 |
| *Coprinellus micaceus* | 20 | 26.8 | 26.5 | 27 | 29.7 | 29.6 | 29.8 | 2.9 |
| *Lactifluus* spp. | 21 | 23.9 (23.7) | 23.9 (23.6) | 24 (23.9) | 25.4 | 25.3 | 25.5 | 1.5 |
| unknown | 22 | 23.7 | 23.7 | 23.8 | 27.9 | 27.8 | 28 | 4.2 |
| unknown | 23 | 23.7 (23.9) | 23.5 (23.7) | 23.8 (24.1) | 25.4 | 25.3 | 25.5 | 1.7 |
| unknown | 24 | 24.9 | 24.7 | 25 | 26.8 | 26.5 | 27 | 1.9 |
| unknown | 25 | 22.6 | 22.5 | 22.7 | 26 | 26.9 | 25.7 | 3.4 |
| unknown | 26 | 25.6 | 25.6 | 25.7 | 27.4 | 27.1 | 27.9 | 1.8 |
| *Boletus spp.* | 27 | 28.5 | 29.1 | 30 | 26.2 | 25.9 | 26.5 | 2.3 |
| *average* |  | 27 (24.3) | 26.9 (24.2) | 28.6 (24.4) | 29.5 | 29.4 | 29.5 | 2.5 |
| *Standard deviation* |  | 4.5 (0.4) | 4.6 (0.4) | 5.1 (0.4) | 4.6 | 4.7 | 4.4 | 1.4 |

**Table S2.** Temperatures (average °C, max/min) of light and dark *A. bisporous* mushrooms caps in normal versus dehydrated states following 1 h incubation at 4, 24, and 37 °C.

| Ambient Temp | Light | | | | | | Dark | | | | | |
| --- | --- | --- | --- | --- | --- | --- | --- | --- | --- | --- | --- | --- |
|  | Normal | | | Dehydrated | | | Normal | | | Dehydrated | | |
|  | avg | max | min | avg | max | min | avg | max | min | avg | max | min |
| 4 | 1.7 | 2.2 | 1.4 | 6.9 | 8.0 | 6.1 | 1.6 | 2.7 | 1.1 | 6.4 | 7.2 | 5.6 |
| 24 | 18.8 | 20.0 | 17.9 | 20.1 | 21.7 | 18.9 | 18.1 | 19.2 | 17.6 | 21.7 | 22.5 | 21.0 |
| 37 | 26.3 | 27.3 | 24.7 | 32.9 | 33.3 | 32.0 | 26.4 | 27.6 | 24.8 | 33.3 | 33.8 | 31.5 |

**Table S3.** Water mass percentage of *A. bisporous* mushrooms caps and *C. neoformans* yeast colonies. Percent water mass was calculated by the mass difference before and after lyophilization. Values represent different biological replicas.

| **Specimen** | **Percent of water (% m/m)** |
| --- | --- |
| *A. bisporous* mushroom caps | 93.6 ± 0.4* |
| Encapsulated *C. neoformans* | 90.2, 89.7, 89.0, 89.6 |
| Acapsular *C. neoformans* | 81.8, 81.8, 82.3, 81.5 |
| Unidentified specimens | 83.3, 88.1, 89.9, 91.6, 82.1, 80.7, 57.4 |

*average and standard deviation from three light and three darkly pigmented specimens.

**Table S4.** Total water mass condensed on lids divided by biofilm or agar area.

| sample | Water mass per area (mg/m^2^) | |
| --- | --- | --- |
|  | biofilm | agar |
| 1 | 6.4 | 0.2 |
| 2 | 10.0 | 0.8 |
| 3 | 5.0 | 0.4 |
| 4 | 3.6 | 0.4 |

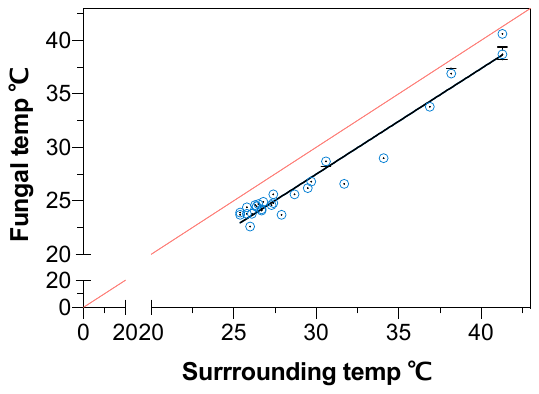

**Fig S1.** Linear regression analysis of yeast, mold, and mushroom cap temperatures as a function of ambient temperature. Slope 0.988, R2 0.95, Y-intercept -2.14 °C, X-intercept 2.16 °C. Red line shows line of identity.

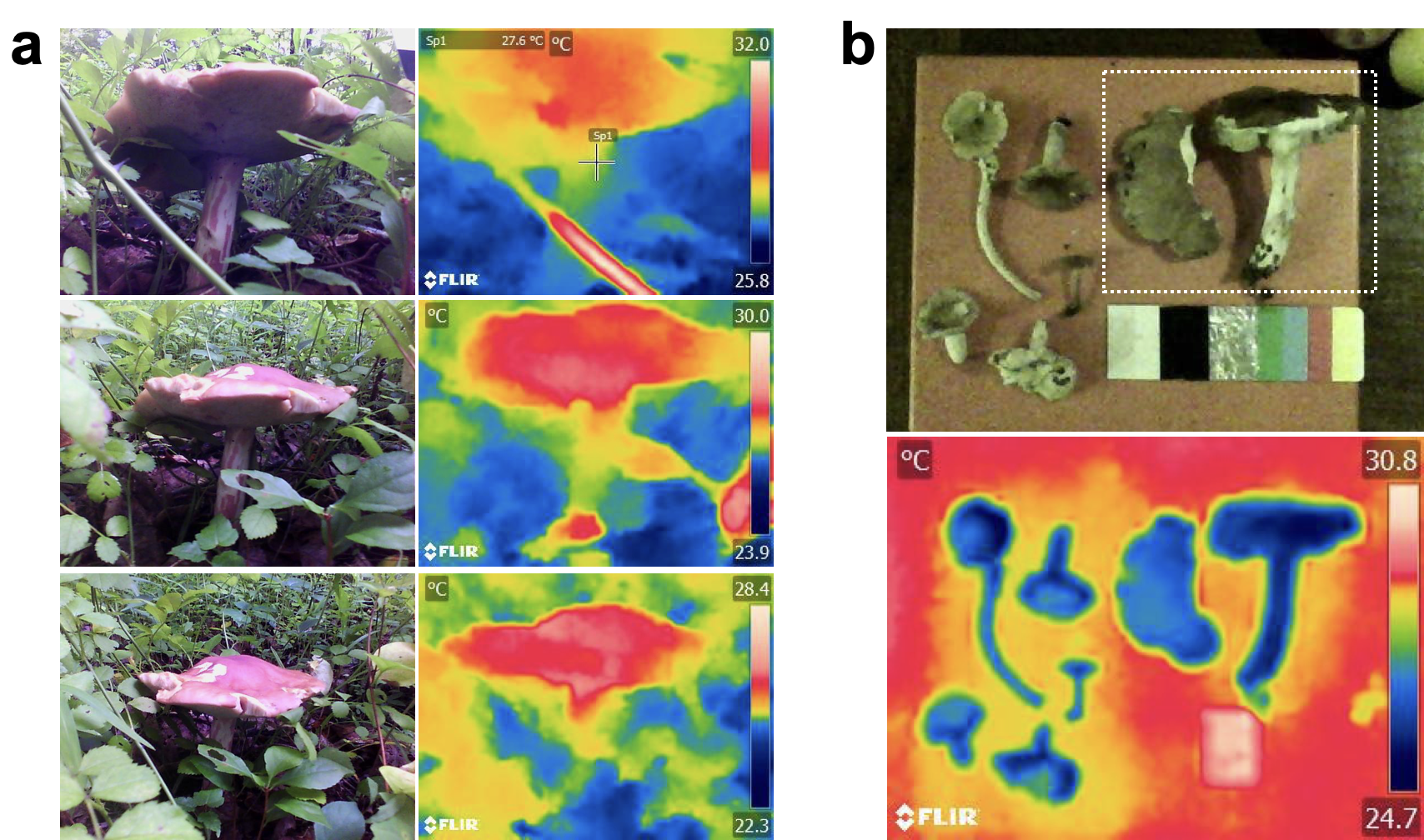

**Fig S2.** (**a**) Visible and infrared images of a *Boletus* spp. mushroom in open-habitat conditions that appear warmer than its surroundings and was exposed to direct sunlight. (**b**) Visible and IR images of six wild-type mushrooms specimens, including the same Boletus mushrooms (inside square), taken indoor, at room temperature, ~4 hours following collection from their natural habitat. Note that all mushroom specimens are colder than surrounding temperatures. See SFig 7 for information about the reference card and vinyl tape as a reference material for ambient temperature.

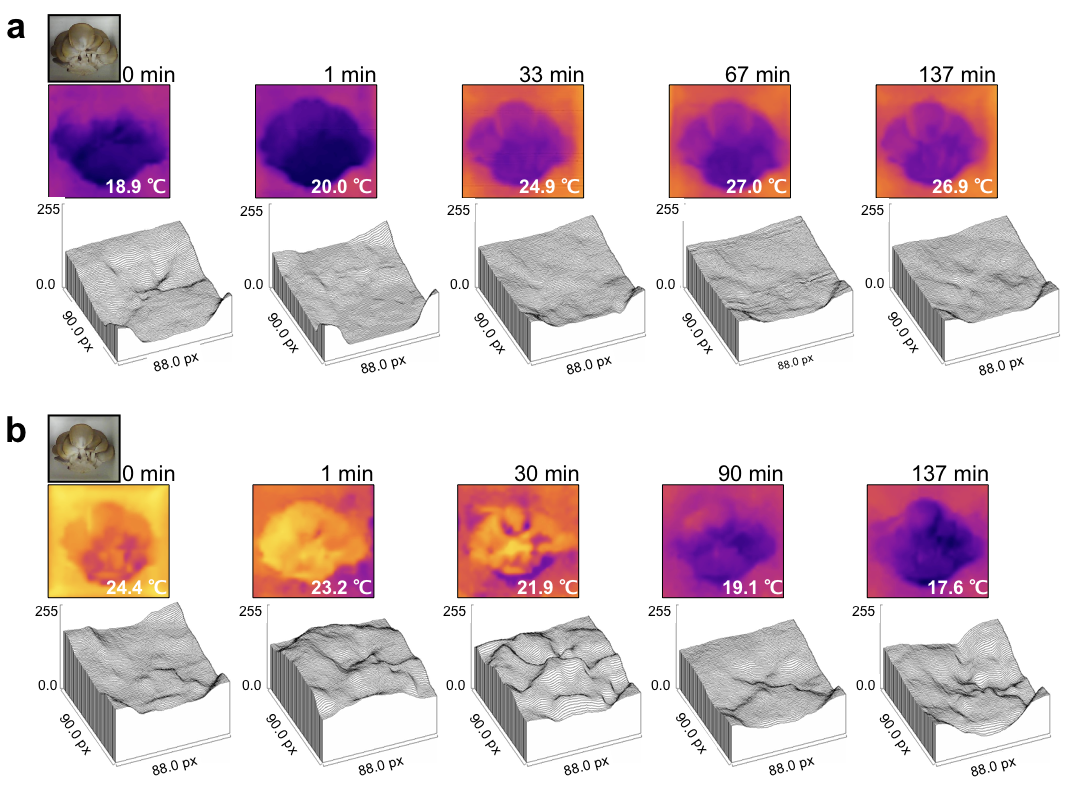
**Fig S3.** Plot profiles of thermal images of *P. ostreatus* mushroom flush detached and subsequently incubated inside a warm room (**a**) at 37 °C (and <10% RH) and a cold room (**b**) 4 °C (and ~30% RH). Inset temperature values correspond to the lowest and highest temperature signal detected in the thermographs in (**a**) and (**b**), respectively.

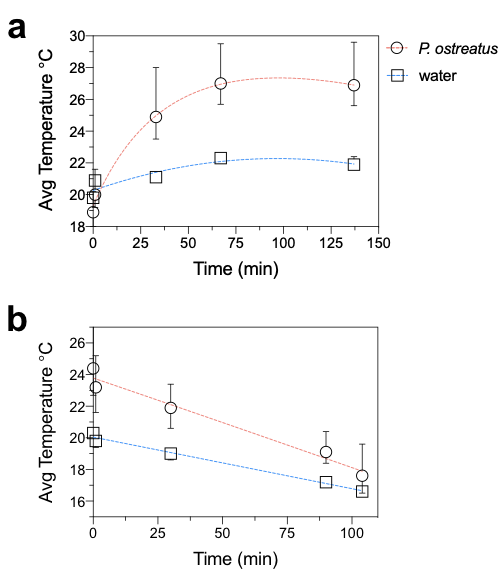

**Fig S4.** *P. ostreatus* mushroom flush (detached) average temperature following incubation at 37 and 4 °C, (**a**) and (**b**), respectively. An equivalent mass of pure water was co-incubated and used as reference.

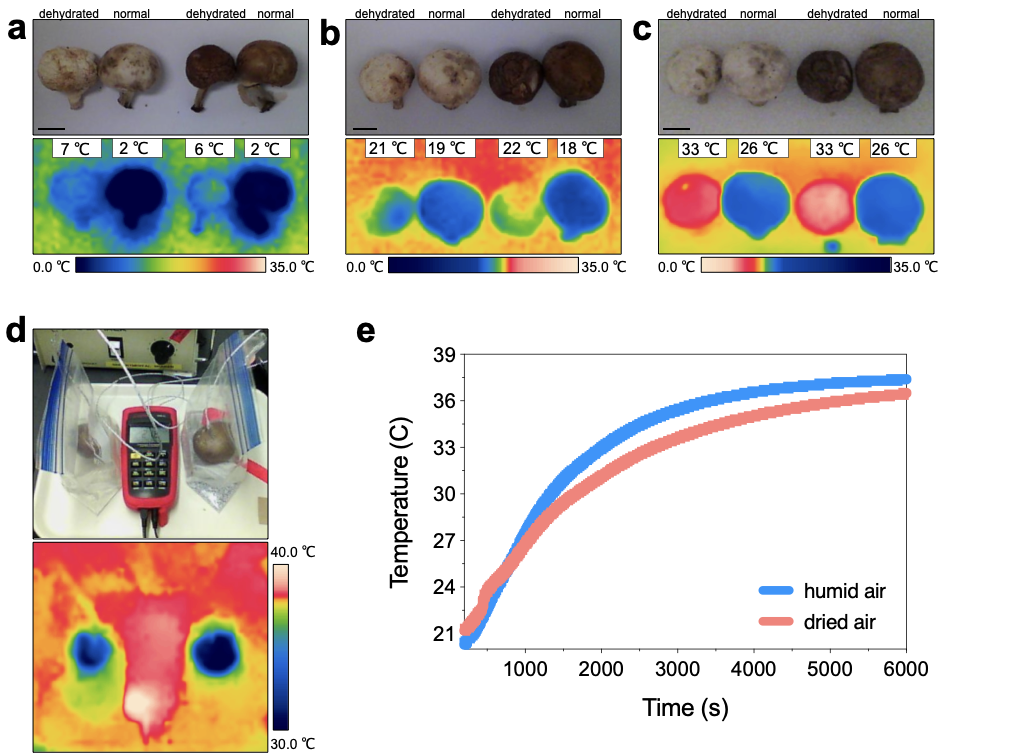

**Fig S5. Mushroom coldness is mediated via evaporative cooling.** Dehydrated (*left*) and normal (*right*) version of white and brown *Agaricus bisporus* detached mushrooms were incubated at: (**a**) 4, (**b**) 23, and (**c**) 37 ºC ambient temperatures. Normal mushrooms are able to maintain cooler temperatures regardless of the ambient temperature. (**d**) Experimental setup to monitor mushroom temperature change as a function of humidity. (**e**) Mushroom temperature as a function of time was recorded inside plastic bags closed (*top*) and opened (*bottom*).

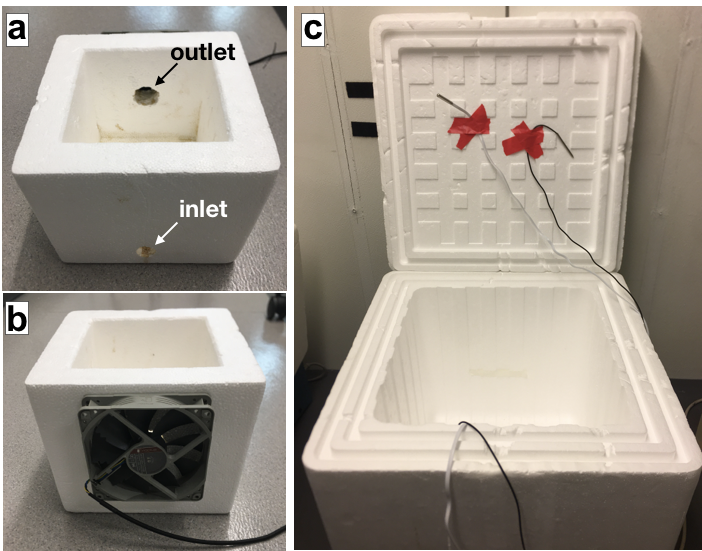
**Fig S6. MycoCooler™ prototype.** (**a**) Large Styrofoam box ((30.48 x 30.48 x 30.48 cm) having an area of 28,317 cm^3^. Temperature and humidity probes tapped to the inner side of the lid can be observed. (**b**) The MycoCooler™ prototype made of a smaller Styrofoam box (lid not shown) with an inlet aperture of 1 cm diameter and outlet aperture of 2 cm diameter. An exhaust fan was glued outside the box centered on top of the outlet aperture.

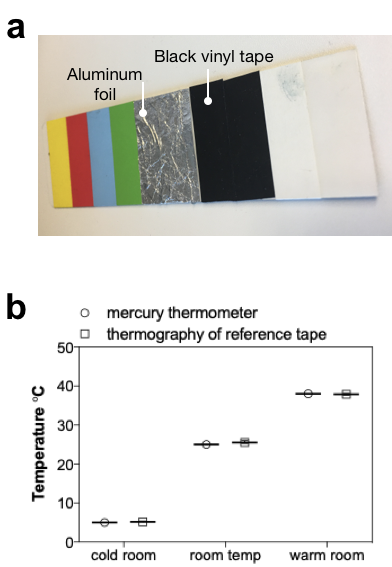

**Fig S7.** **Black vinyl tape as reference for ambient temperature.** (**a**) Reference card containing black vinyl tape and aluminum foil. (**b**) Temperatures measurements inside a cold (5 °C), warm (37 °C), and regular (25 °C) rooms using a mercury thermometer and thermography of reference card-black tape.
